# Germ Granule Remodeling Reveals Germline Age in *Caenorhabditis elegans*

**DOI:** 10.64898/2026.11.10.765724

**Authors:** Purbasha Dasgupta, Ryan J. Lee, Erica Dinneen, Carlos G. Silva-García

## Abstract

Germline aging limits reproductive lifespan, yet quantitative measures of germline age at cellular resolution within intact tissue remain limited. Here, we show that two germline condensates, P granules and Mutator foci, undergo systematic age-associated remodeling in *Caenorhabditis elegans*, changing in morphology, spatial organization, and mutual associations. From these single-nucleus features, we develop Germ Granule Clock (GG-Clock), an image-based aging clock that accurately predicts chronological age. Intermittent fasting, a pro-longevity intervention, shifts GG-Clock predictions toward a younger germline state, demonstrating the clock’s responsiveness to anti-aging interventions. To make the approach broadly usable, we introduce GGProfiler, an open-source tool that automates granule segmentation and feature extraction. Together, our findings establish germ granule architecture as an interpretable, image-based readout of germline aging and provide a platform for quantifying condensate remodeling across genetic and environmental interventions, with potential translation to other systems.

## MAIN TEXT

Robust biomarkers of germline aging are needed to understand reproductive decline and evaluate interventions. Such biomarkers can underpin aging clocks that quantitatively estimate germline age. Although aging clocks now span multiple molecular and imaging modalities (Liu et al., 2026), few target the germline, and somatic clocks may not capture germ cell aging (Horvath, 2013). Existing germline clocks, including epigenetic clocks for mammalian gametes (Jenkins et al., 2018; Kordowitzki et al., 2021; Pilsner et al., 2022) and a single-nucleus transcriptomic clock in *C. elegans* (Gao et al., 2024), require destructive sequencing workflows. These workflows lose native spatial context and preclude repeated measurements of the same cells. Imaging offers a complementary approach, with germ granules serving as accessible targets. These conserved, phase-separated ribonucleoprotein condensates mediate RNA surveillance, small-RNA amplification, and transposon silencing (Voronina et al., 2011), and their organization remodels during development and under stress (Tanaka et al., 2011; Uebel et al., 2023), suggesting that germ granules report on the physiological state of the germline. Whether their architecture also reflects aging is unknown. We focused on P granules and Mutator foci, constitutive perinuclear condensates present in the unstressed *C. elegans* germline, rather than stress-induced ribonucleoprotein (RNP) granules (Schisa, 2014; Huelgas-Morales et al., 2016). We hypothesized that their morphology, number, and spatial organization could provide quantitative readouts of germline aging. Because *C. elegans* is optically transparent, these condensates can be imaged in the intact germline of live animals, making it possible to build an interpretable, image-based germline aging clock.

To test this hypothesis, we imaged germ granules surrounding pachytene nuclei in the germline of live, anesthetized *C. elegans* hermaphrodites across adult days 1–5 in two independent cohorts (Figure 1A). Restricting sampling to pachytene nuclei provides a defined meiotic context and limits developmental variation, particularly granule redistribution during oocyte differentiation (Pitt et al., 2000; Sheth et al., 2010). We visualized two compartments: P granules (PGL-1::mCardinal) and Mutator foci (MUT-16::GFP). These neighboring compartments have connected roles in RNA surveillance and small RNA amplification but respond differently to perturbation (Uebel et al., 2023). Thus, quantifying their individual architecture and spatial association allowed us to assess age-associated changes within and between them. Using standard two-channel fluorescence microscopy, we acquired z-stacks and selected paired nuclear crops from the same focal plane in both channels (Supplementary File 1). This avoids superimposing granules from different depths, which could merge distinct objects or create apparent proximity in projected images (Dunn et al., 2011; Marnik et al., 2022). Independent segmentation and feature extraction yielded twelve measurements describing condensate number, size, shape, spacing, radial organization, and association within the sampled plane.

**Figure 1.**
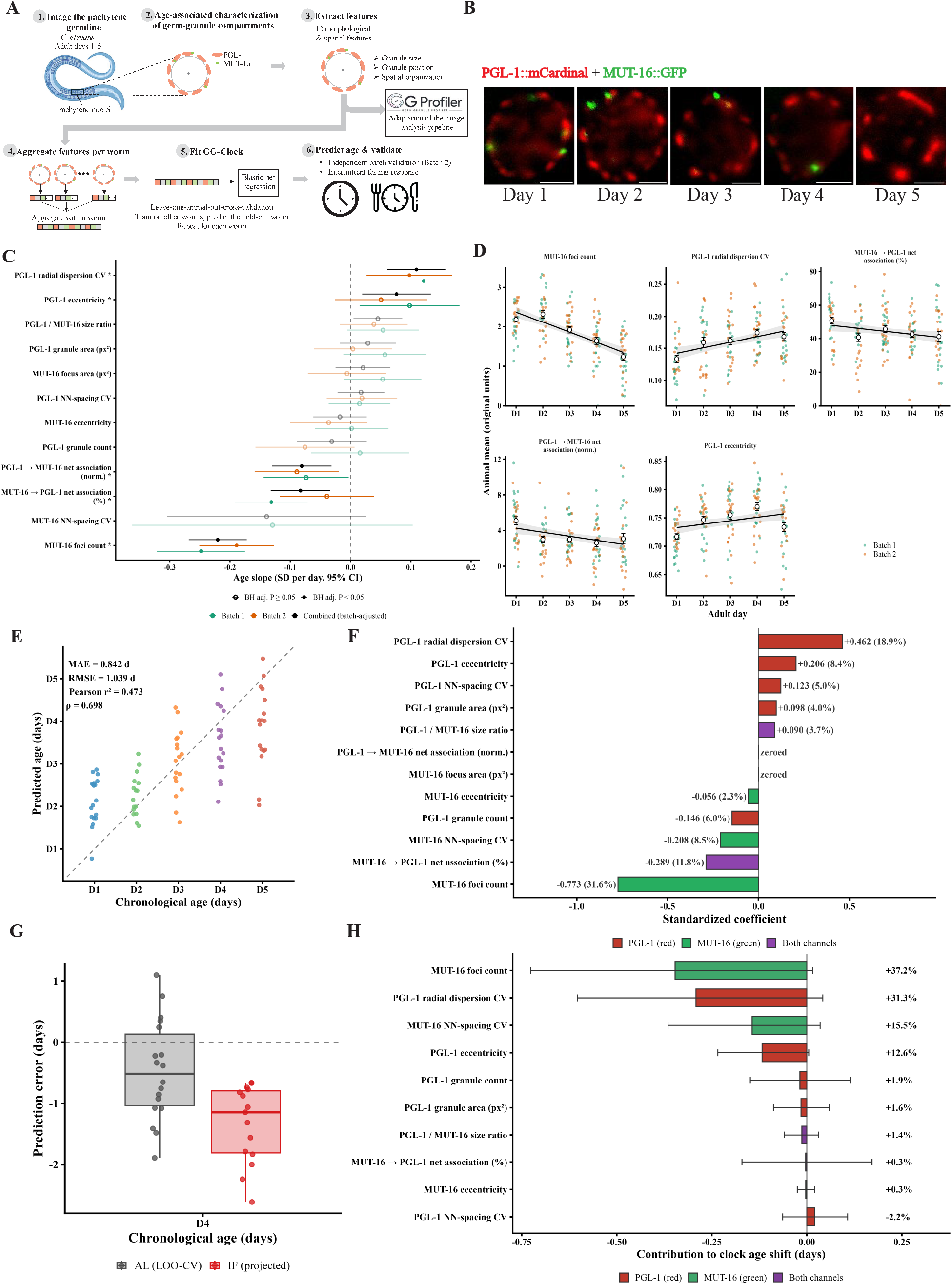
Germ-granule architecture remodels with age and enables an aging clock (GG-Clock). **(A)** Study workflow. Pachytene germline nuclei of *C. elegans* were imaged at adult days 1–5, with PGL-1::mCardinal (red) and MUT-16::GFP (green) marking two germ-granule compartments (1-2). Twelve morphological and spatial features were extracted and aggregated per worm (3-4). An elastic-net regression (GG-Clock) was trained to predict age (5-6). It was validated by leave-one-out cross-validation (LOO-CV) and tested for responsiveness in intermittent-fasting (IF) animals. The image-analysis workflow was adapted into GGProfiler. **(B)** Representative images showing age-associated germ-granule remodeling. [Scale bar, 2 µm.] **(C)** Age-associated changes in germ-granule architecture. Points show the age slope (SD per day) with 95% CI for each of the 12 features, from linear mixed-effects models with nuclei nested within animals. Results are shown for Batch 1 (green; up to 777 nuclei, 89 animals), Batch 2 (orange; up to 807 nuclei, 89 animals), and both batches combined (black; batch-adjusted). Asterisks mark features with a combined BH-adjusted P < 0.05. Faded features show no combined age effect. No feature showed a batch × age interaction (all BH-adjusted P ≥ 0.62). **(D)** Trajectories of the five features with a significant combined age effect. Points are individual animals (Batch 1 and Batch 2); open circles are day means ± SE of animal means. Lines and bands show the linear fit with 95% CI. Slopes and BH-adjusted P values come from the combined, batch-adjusted nested mixed model (178 animals). **(E)** GG-Clock accuracy under leave-one-out cross-validation (LOO-CV). Each point is one ad libitum-fed (AL) animal (n = 89; colored by chronological age), and the dashed line marks perfect prediction. MAE = 0.842 d, RMSE = 1.039 d, Pearson r^2^ = 0.473, Spearman ρ = 0.698. **(F)** Elastic-net feature weights of GG-Clock, shown as standardized coefficients (coefficient × training SD). Percentages give each feature’s share of total absolute weight; features with zero weight are labeled “zeroed”. Bars are colored by channel: PGL-1 (red), MUT-16 (green), or both channels (purple). **(G)** Clock prediction error (predicted − chronological age) at adult day 4 for ad libitum-fed (AL; gray; LOO-CV predictions, n = 18) and intermittently fasted (IF; red; n = 15) animals (Batch 1). IF animals were projected onto the AL-trained clock. IF animals were assigned a younger age than AL animals (−0.87 d, 95% CI −1.40 to −0.35; one-sided p = 0.001, linear model). Boxes show the median and IQR; whiskers extend to 1.5 × IQR. **(H)** Feature contributions to the day-4 IF-versus-AL shift in predicted age (Batch 1; n = 18 AL, 15 IF; clock weights fixed; negative = younger). Mutator-focus number and P-granule radial variability together accounted for 68% of the shift (95% CI 32–101%). Whiskers, 95% bootstrap CI (5,000 resamples of animals). Colors as in panel F. Statistical details for panels C–D are provided in Supplementary Tables S1–S4, and those for panels E–H in Supplementary Tables S5–S11.

Germ granule architecture remodels with age (Figure 1B). Five of twelve features show significant linear trends with age in a combined analysis of the two cohorts (Figure 1C-D). The greatest change is a decline in Mutator-focus number. Among images with both compartments present, Mutator foci and P granules show reduced above-chance proximity to each other. P granules are also more elongated and more variable in their distances from the nuclear center, consistent with altered spatial organization. The remaining features do not change significantly (Figure 1C). Together, these changes suggest compartment-specific remodeling, with aging affecting Mutator-focus abundance and P-granule shape and spatial organization. Our data on these distinct age-associated responses parallel previous observations that Mutator foci and P granules differ in their sensitivity to perturbation (Uebel et al., 2023).

We next asked whether this remodeling contains sufficient information to predict chronological age. For this, we developed GG-Clock (Germ Granule Clock), an elastic net-based predictor trained on animal-level profiles aggregated from nucleus-level measurements in *ad libitum*-fed animals at adult days 1–5 (Figure 1A). This approach retains the animal as the unit of biological replication. In leave-one-animal-out cross-validation, GG-Clock achieves a mean absolute error (MAE) of 0.842 days (Spearman ρ = 0.698; Figure 1E). We then applied the model without retraining to an independent cohort, where it performed comparably (MAE = 0.858 days; Spearman ρ = 0.689; Supplementary Figure 1A), supporting reproducible age prediction across independent cohorts. Standardized feature weights identify Mutator-focus number as the strongest contributor, followed by variability in P-granule distance from the nuclear center, above-chance proximity of Mutator foci to P granules, Mutator-focus spacing variability, and P-granule elongation (Figure 1F). Higher Mutator-focus number and proximity contribute to younger predicted ages, whereas greater radial variability and elongation of P granules contribute to older predicted ages. Four of these five features (all except Mutator focus spacing variability) also show significant age-associated trends in the same direction (Figure 1C-D), linking the clock’s predictions to the observed architectural remodeling. To our knowledge, GG-Clock is therefore the first imaging-based predictor of germline age, built directly from age-responsive features of germ-granule architecture.

We next tested whether GG-Clock responds to intermittent fasting (IF), a regimen that extends lifespan in *C. elegans* (Dasgupta & Silva-García, 2026). At adult day 4, the clock assigns IF animals a younger age than *ad libitum* controls (−0.87 d, 95% CI −1.40 to −0.35; p = 0.001; Figure 1G). Feature-contribution analysis attributes much of the younger prediction to higher Mutator-focus number and lower P-granule radial variability, both opposite to their age-associated trajectories (Figure 1H). This response is further consistent across independent batches (Supplementary Figure 1B-C). Thus, GG-Clock responds to a pro-longevity intervention, shifting the germ-granule signature toward a younger state in animals of the same chronological age.

Germ granule architecture continues to change after the reproductive period. In a parallel subset of unmated hermaphrodites drawn from the same population, daily progeny peaks at day 2 and falls to zero by day 6 (negative binomial mixed model, likelihood-ratio test p < 0.001, Supplementary Figure 2A), with animals having produced, on average, 97.3% of their total brood by day 4 (n = 15, Supplementary Figure 2B). To ask whether germline age keeps advancing beyond this point, we retrained the GG-Clock on days 1–6. The extended clock performs equivalently on days 1–5 (MAE 0.848 d vs 0.842 d), so we retained the days 1–5 clock for all main analyses and used the days 1–6 clock only for this comparison (Supplementary Figure 2C, Table S26). In this model, predicted age increases by 1.35 days between days 4 and 6 (95% bootstrap CI 0.73–1.99; p < 0.001; Supplementary Figure 2D). Granule remodeling therefore continues after progeny production is largely complete, suggesting that GG-Clock tracks an ongoing age-associated process rather than cumulative reproductive output. However, because sperm depletion can remodel ribonucleoprotein granules both in the pachytene region and in arrested oocytes (Schisa, 2014), the rise predicted after day 4 could partly reflect exhaustion of self-sperm. Whether male sperm provided by mating, which extends the reproductive period, alters this trajectory remains to be tested, although such experiments are complicated by the variable number of sperm transferred during mating.

Finally, germ-granule quantification has relied largely on fixed, dissected germlines, manual analysis, or study-specific pipelines (Toraason et al., 2021; Uebel et al., 2023). To make our image-analysis framework broadly accessible, we built GGProfiler (v1.0), an open-source, code-free Shiny application (Figure 2A) that automates segmentation, feature extraction, statistical comparison, and visualization of germ granule architecture (Figure 2B). As a demonstration, we profiled germ-granule architecture after acute heat shock (32 °C for 2 h) and starvation (8 h). Heat shock causes broad remodeling (Figure 2C-E, Table S24). PGL-1 granules become fewer, smaller, and less elongated; MUT-16 foci are depleted and more elongated; and PGL-1 and MUT-16 overlap is markedly reduced, consistent with the heat shock sensitivity of the Mutator foci reported previously (Uebel et al., 2023). Starvation has more limited effects, reducing the number of both PGL-1 granules and MUT-16 foci, with modest changes in PGL-1 elongation and PGL-1/MUT-16 overlap (Figure 2F-H, Table S25). Full methods and feature definitions (Table SF1) are provided in the Materials and Methods section in Supplementary File 1. GGProfiler thus makes germ-granule architecture an accessible phenotype for studying germline physiology across biological contexts.

**Figure 2.**
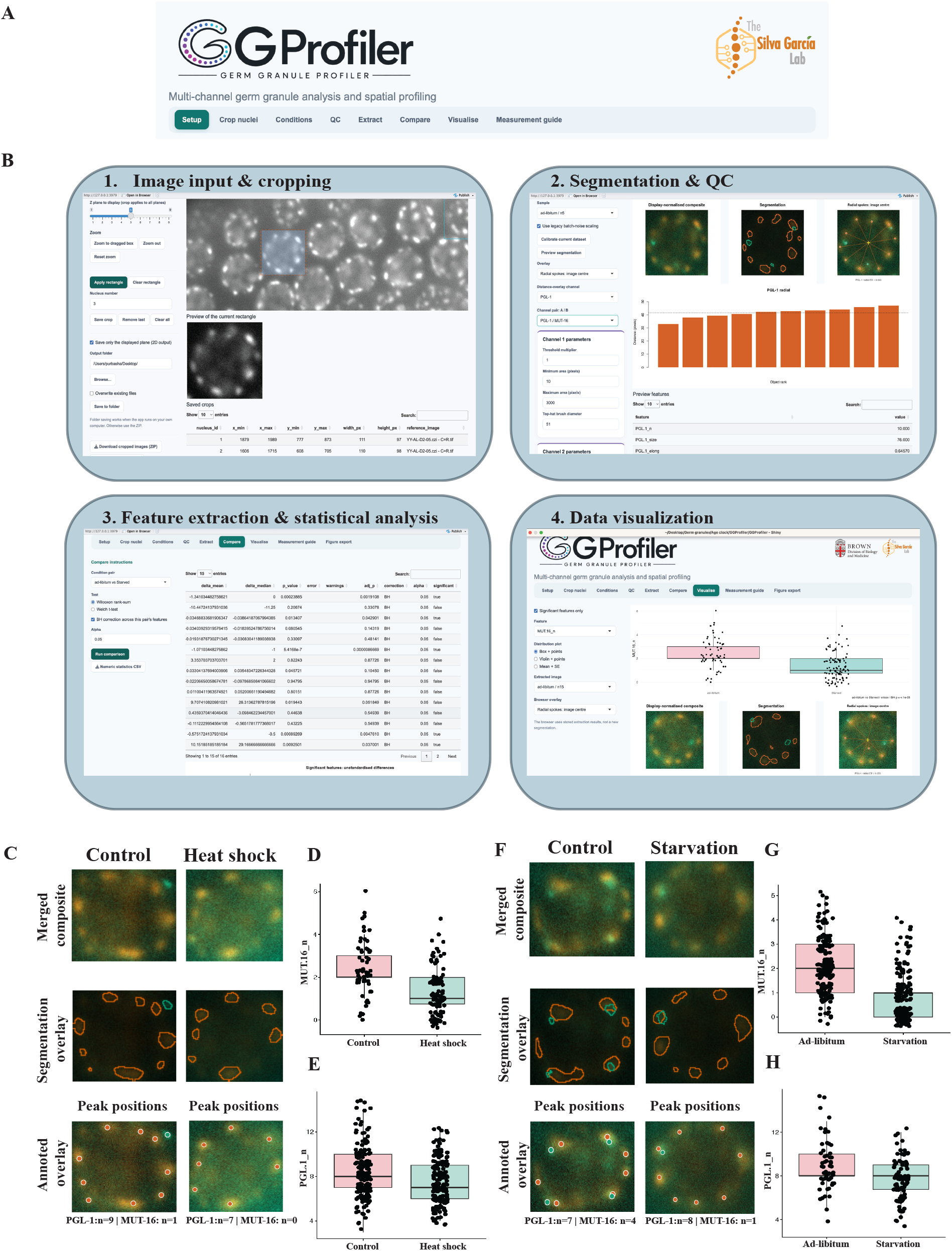
GGProfiler, an open-source, code-free platform for profiling germ-granule architecture, reveals stress-induced changes. **(A)** The GGProfiler interface, with modules for Setup, Crop nuclei, Conditions, QC, Extract, Compare, Visualize, Measurement guide, and Figure export. **(B)** The four-step workflow: (1) image input and nuclear cropping, (2) segmentation and QC, (3) feature extraction and statistical analysis, and (4) data visualization. **(C)** GGProfiler-generated outputs for a representative pachytene nucleus from control and heat-shocked animals: merged composite, segmentation overlay, and annotated overlay with detected peak positions. PGL-1 and MUT-16 granule counts for the nucleus shown are given below the images. Heat shock at 32 °C for 2 h. **(D, E)** GGProfiler-generated quantification of MUT-16 foci (D; box plot, points = nuclei) and PGL-1 granules (E; box plot, points = nuclei) in control and heat-shocked animals. Wilcoxon test, BH-adjusted P < 1 × 10^-5^ (D) and P = 2.1 × 10^-4^ (E). **(F)** As in (C), for ad libitum-fed and starved animals. Starvation duration: 8 h. **(G, H)** As in (D, E), for ad libitum-fed versus starved animals. Wilcoxon test, BH-adjusted P < 1 × 10^-5^ (G) and P = 0.0019 (H). GGProfiler generated statistical details for the heat-shock comparisons (D, E) are in Table S24, and for the starvation comparisons (G, H) in Table S25.

In conclusion, our study characterizes age-associated remodeling of two germ-granule compartments, P granules and Mutator foci, in terms of their morphology, spatial organization, and abundance, providing, to our knowledge, the first quantitative description of age-associated remodeling of these condensates. Building on these changes, we develop GG-Clock, the first imaging-based, germline-specific aging clock derived from condensate architecture. It provides a minimally invasive, imaging-based approach to age prediction without tissue dissociation or sequencing. Cross-cohort validation and responsiveness to intermittent fasting support its robustness, and GGProfiler makes the framework accessible to the broader community. Future work should evaluate both tools across reproductive contexts, genetic backgrounds, imaging platforms, and additional germ granule compartments. More broadly, the conservation of germ granules raises the question of whether condensate remodeling is a shared feature of germline aging across species and whether it could serve as an indicator of reproductive health.

## MATERIALS AND METHODS

Materials and Methods can be found in Supplementary File 1. The authors affirm that all data necessary for confirming the conclusions of the article are present within the article, figures, and tables included in the Main Text File, Supplementary File 1, and Supplementary File 2.

## Supporting information

GGProfiler v1.0 user manual

## DATA AVAILABILITY STATEMENT

The source code for GGProfiler (v1.0) and analysis scripts are available on GitHub at https://github.com/Purbashadg/Germ-Granule-Remodeling. Raw microscopy images are available from the corresponding author upon request. Supplementary Tables S1–S26 provide detailed statistical results and are included in Supplementary File 2.

## ACKNOWLEDGEMENTS

We thank all members of the Silva-García laboratory for their helpful discussions, and Sean Ovalles in particular for his comments on the manuscript and data. The YY1492 *[mut-16(cmp3[mut-16::gfp::flag + loxP]) I; znfx-1(gg634[HA::tagRFP::znfx-1]) II; pgl-1(gg640[pgl-1::3xflag::mCardinal]) IV]* strain was provided by the Caenorhabditis Genetics Center (CGC), which is funded by the NIH Office of Research Infrastructure Programs (P40 OD010440).

## AUTHOR CONTRIBUTIONS

P.D. and C.G.S.G. designed the study. P.D. processed images, conducted the statistical analyses, and developed GG-Clock and GGProfiler. R.J.L. acquired all microscopy images and contributed to image processing. E.D. contributed to image processing. P.D. and C.G.S.G. wrote the manuscript, incorporating comments and revisions from all authors.

## EXTENDED DATA

**Supplementary Figure 1.**
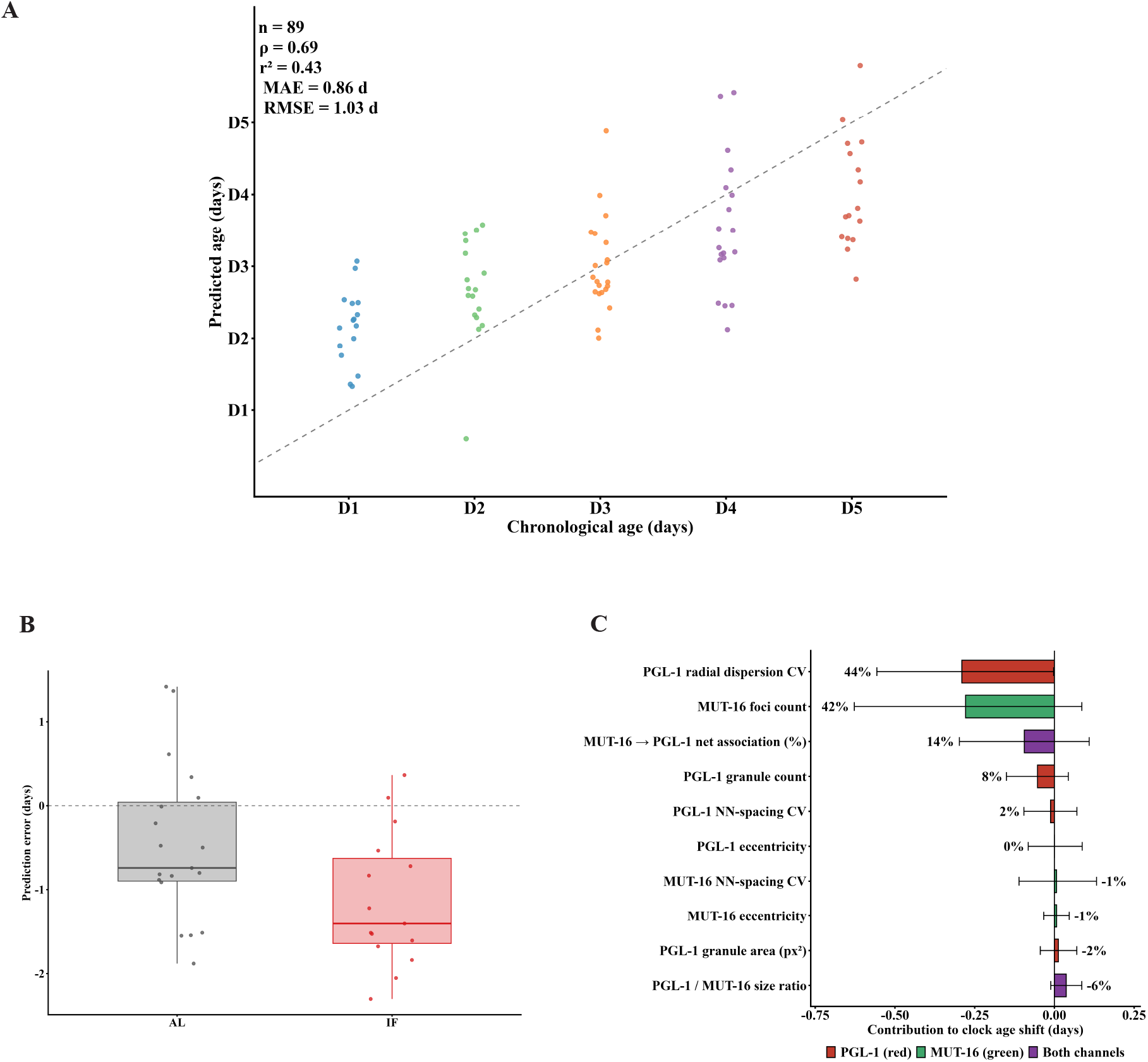
GG-Clock generalizes across experimental batches. **(A)** External validation of the Batch 1 clock. Predicted versus chronological age for Batch 2 ad libitum-fed (AL) animals (n = 89), scored with the Batch 1-trained clock and the Batch 1 image-noise reference. The clock was not refit or tuned on Batch 2. The dashed line marks perfect prediction. Spearman ρ = 0.69, Pearson r^2^ = 0.43, MAE = 0.86 d, RMSE = 1.03 d. **(B)** Prediction error at adult day 4 for Batch 2 ad libitum-fed (AL; gray; external, n = 19) and intermittent-fasting (IF; red; projected, n = 15) animals scored with the Batch 1 clock. IF animals were assigned a younger age than AL animals (−0.67 d, 95% CI −1.28 to −0.05; one-sided p = 0.017, linear model). Boxes show the median and IQR; whiskers extend to 1.5 × IQR. **(C)** Feature contributions to the younger clock age of Batch 2 IF animals at day 4 (n = 19 AL, 15 IF), with Batch 1 clock weights held fixed. Percentages show each feature’s share of the total shift (−0.67 d, 95% CI −1.25 to −0.09); negative shares oppose the shift. Whiskers show 95% bootstrap CIs over animals (5,000 resamples). Two features with zero clock weight are omitted. Bars are colored by channel as in Figure 1F. Statistical details are provided in Supplementary Tables S12–S17.

**Supplementary Figure 2.**
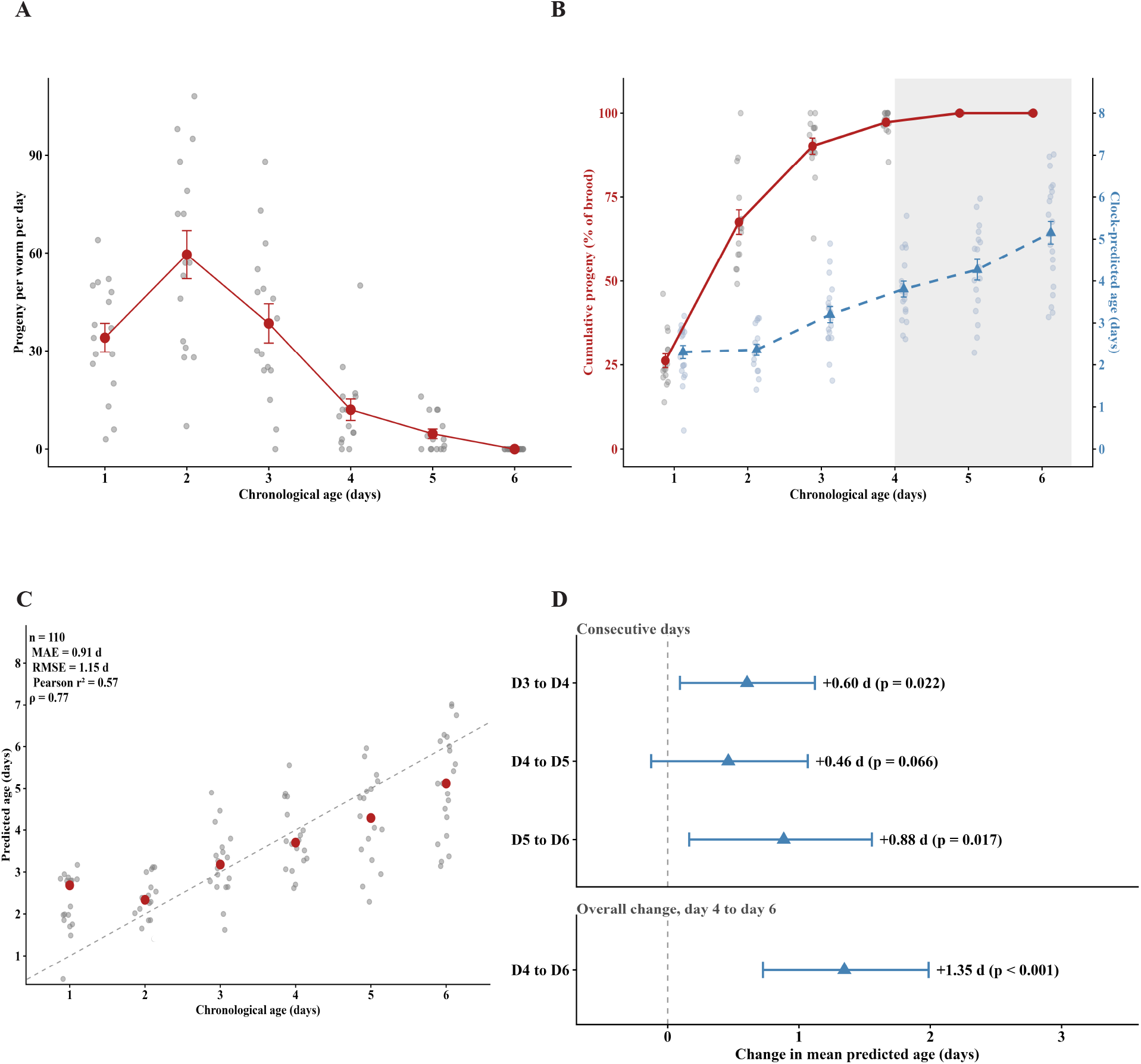
GG-Clock-predicted age continues to increase after most reproductive output has occurred. **(A)** Daily progeny per hermaphrodite across adult days 1–6. Gray points are individual animals, and red symbols are mean ± SEM. Daily progeny varied with age (negative binomial mixed model, likelihood-ratio test p < 0.001, days 1–5; day 6 was excluded because all counts were zero). n = 15 self-fertile hermaphrodites followed daily. **(B)** Cumulative progeny and clock-predicted age across adult days 1–6. Red (left axis) shows cumulative progeny as a percentage of each animal’s total measured brood. Gray points are individual animals, and dark red symbols are mean ± SEM (n = 15 animals with complete records). Blue (right axis) shows predicted age from the GG-Clock trained on days 1–6 (leave-one-out cross-validation). Light blue points are individual animals, and dark blue symbols are mean ± SEM (n = 110). Shading marks day 4 onward, when mean cumulative output reached ≥ 95% of the total brood.Progeny and clock data come from separate cohorts. **(C)** Predicted versus chronological age for days 1– 6 GG-Clock (leave-one-out cross-validation, n = 110 animals). Gray points are individual animals, and red circles are mean ± SEM. The dashed line marks predicted age equal to chronological age. Overall MAE = 0.909 d, RMSE = 1.147 d, Pearson r^2^ = 0.569 and Spearman ρ = 0.772. On days 1–5 (n = 89 animals), the days 1–6 clock had an MAE of 0.848 d, compared with 0.842 d for the days 1–5 clock used in the main analysis. **(D)** Change in mean predicted age between days for the days 1–6 clock, calculated as the mean predicted age on the older chronological day minus that on the younger chronological day. Points show the difference in means, and bars show 95% bootstrap confidence intervals (5,000 resamples of animals within each day). The upper panel shows consecutive days, and the boxed lower panel shows the overall change from day 4 to day 6 (+1.35 d, 95% CI 0.73–1.99). P values are from one-sided Mann–Whitney U tests. Statistical details are provided in Supplementary Tables S18–S23 & S26.

