## Supplementary File 1 for "Germ Granule Remodeling Reveals Germline Age in *Caenorhabditis elegans*"

### MATERIALS AND METHODS

#### *C. elegans* strains and husbandry

The strain used in this study, YY1492 [mut-16(cmp3[mut-16::gfp::flag + loxP]) I; znfx-1(gg634[HA::tagRFP::znfx-1]) II; pgl-1(gg640[pgl-1::3xflag::mCardinal]) IV] (Wan et al., 2018), was obtained from the Caenorhabditis Genetics Center, which is funded by the NIH Office of Research Infrastructure Programs (P40 OD010440). Worms were maintained at 20 °C on standard nematode growth medium (NGM) seeded with *E. coli* OP50-1.

#### Age synchronization and imaging

Worms were age-synchronized by bleaching and maintained in the absence of FUdR. For imaging, at the indicated adult days, worms were immobilized in 10× diluted 100 mM tetramisole, mounted on 2% agarose pads, and imaged on a Zeiss Axio Imager Z2 with a 63×/1.4 Plan Apo oil objective. Z-stacks were acquired through the gonad in DIC, GFP, and mCardinal channels. tagRFP::ZNFX-1 was excluded from analysis to avoid spectral bleed-through.
For clock construction, about 16–20 worms were imaged per day on adult days 1–5 in two independent biological replicates (Batch 1, discovery cohort; Batch 2, replication cohort; animals and nuclei per condition and day in Tables S10 and S14). For the extended clock, animals were also imaged on day 6.

#### Intermittent fasting and acute stress treatments

For intermittent fasting (IF), worms underwent repeated 5 h fasting cycles daily from adult day 1, following the protocol of Dasgupta & Silva-García (2026). They were imaged on day 4 after overnight refeeding. Untreated controls (about 16–20 worms per condition) were imaged alongside the IF worms in both replicates. To test the sensitivity of GGProfiler to acute perturbations, about 10–20 synchronized day-1 worms were starved for 8 h or heat-shocked at 32 °C for 2 h. They were imaged immediately afterward alongside controls, using the same imaging protocol.

#### Image processing and segmentation

From the z-stacks acquired for each worm, split-channel image stacks (PGL-1::mCardinal, red; MUT-16::GFP, green) were loaded into Fiji (ImageJ) (Schindelin et al., 2012), and paired single-nucleus crops were generated using a custom macro. For each nucleus, a rectangular region of interest (ROI) was manually drawn on the red channel (PGL-1::mCardinal) at its best-focus z-slice. The ROI was fitted tightly along the periphery of the nucleus, so the center of each cropped image approximated the nuclear center. We then applied the same rectangle and z-slice to the other channel, duplicated each channel as a single-plane image, and saved it as an independent grayscale TIFF with paired filenames encoding the nucleus index. The macro validated that the ROI was applied identically to both channels, confirmed that each duplicate and saved file was a single-plane grayscale image of the expected dimensions, and re-opened saved files to verify on-disk integrity before proceeding, halting on any pair that failed validation. This yielded 6–10 paired single-nucleus, single-channel TIFF images per worm for downstream analysis without modifying the original multi-channel z-stacks.

All downstream image processing, segmentation, and feature extraction were performed in R (v4.5.0) using the EBImage package (v4.50.0) (Pau et al., 2010), operating directly on these paired single-channel TIFFs. Red and green channels were processed as independent grayscale matrices at every segmentation and feature-extraction step. They were combined only to generate two-color overlay images for QC and representative purposes. Both biological replicates used for the clock were processed through an identical computational pipeline. The same core pipeline underlies GGProfiler, with additional user-configurable parameters to allow flexible adaptation to different experimental contexts.

P granules marked by PGL-1::mCardinal (red channel) were Gaussian-smoothed ($\sigma = 1$ px), and a local background estimate was obtained by grayscale morphological opening with a 51-pixel disc structuring element (top-hat background subtraction; Bright & Steel, 1987). The background-subtracted residual was thresholded at mean + 1.0 × SD of positive residual pixels. Thresholded masks were hole-filled and morphologically closed (3-pixel disc) prior to connected-component labeling. Labeled objects with area outside 10–3,000 $px^2$ were discarded.

Mutator foci marked by MUT-16::GFP (green channel) were enhanced using a Difference-of-Gaussians bandpass filter ($\sigma_1 = 1$ px; $\sigma_2 = \min(8 \text{ px}, \text{image minimum dimension}/6)$), consistent with a Marr–Hildreth-type blob detector (Marr & Hildreth, 1980), and thresholded at mean + 2.0 × SD of DoG-positive pixels. Masks were hole-filled, morphologically closed (3-pixel disc), and connected-component labeled. Each candidate focus was then required to pass four gates to be retained: (i) an area gate (20–1,000 $px^2$); (ii) a global intensity gate, requiring the focus's mean intensity to exceed the whole-image mean + 1.3 × SD; (iii) an eccentricity gate ($\leq$ 0.90), excluding elongated, streak-like artifacts; and (iv) an adaptive local-contrast gate, requiring the focus's mean intensity to exceed its local background (median intensity within a 4-pixel dilation ring around the object, excluding the object itself) by at least a z-score threshold, computed as (focus mean − local background median) / local background median absolute deviation (MAD). The local-contrast z-threshold (baseline z = 1.0) was scaled per image by that image's median local-background MAD relative to a batch-wide reference (the median across all images in the batch), so that images with intrinsically higher background speckle (e.g., later adulthood days) required proportionally greater local contrast for a focus to be retained. Only foci passing all four retention criteria were kept as final objects. All segmentation and gating parameters were optimized empirically by visual inspection of QC overlay panels spanning a representative set of images across all experimental conditions.

**Feature extraction**

Feature extraction was done in R (v4.5.0) with the EBImage package. Twelve morphological and spatial features were extracted from each nucleus image and aggregated per worm. Counts (PGL-1 granule count, MUT-16 foci count) were averaged across nuclei, and all other features were summarized by the median of finite values. The features described object area (PGL-1 granule area, MUT-16 focus area), shape (PGL-1 and MUT-16 eccentricity), spatial regularity (PGL-1 and MUT-16 NN-spacing CV, PGL-1 radial dispersion CV), cross-channel proximity (MUT-16 → PGL-1 net association (%), PGL-1 → MUT-16 net association (norm.)), and relative size (PGL-1 / MUT-16 size ratio). NN-spacing CV was the coefficient of variation of nearest-neighbor distances between object intensity peaks, and radial dispersion CV was the coefficient of variation of PGL-1 granule peak distances from the image center. Net association indices were the percentage of objects whose intensity peak fell within the other channel's mask (dilated with a 7-px disc), corrected for chance overlap. Images with no detected MUT-16 foci were retained with a foci count of 0, and the MUT-16-dependent features (MUT-16 focus area, MUT-16 eccentricity, MUT-16 → PGL-1 net association) were set to missing for these images before aggregation. Intensity-based features were excluded to reduce sensitivity to differences in laser power, exposure, and photobleaching between sessions. Exact definitions and calculation details for each feature are given below in Table SF1.

**Table SF1. Definitions and interpretation of germ-granule architectural features.**

| Feature | Channel | Formula | Interpretation |
| --- | --- | --- | --- |
| **PGL-1 granule count** | PGL-1 | $$\overline{N_R} = \frac{1}{K}\sum_{k=1}^{K} n_k^R$$ <br> *$n_k^R$= PGL-1 granule counts in nucleus k; K= total nuclei imaged per worm* | Mean P granule count per nucleus |
| **PGL-1 granule area** | PGL-1 | $$\widetilde{a_{R,k}} = \text{median}\left(a_{1,k}^R, \dots, a_{n_k^R,k}^R\right), \quad n_k^R > 0$$ <br> $$A_R = \text{median}\left(\widetilde{a_{R,k}} : k \in V_R\right)$$ <br> *$a_{i,k}^R$ =area of PGL-1 granule (i) in nucleus k;$\widetilde{a_{R,k}}$ = nucleus-level median PGL-1 granule area; $V_R$ = set of nuclei of the animal with finite nucleus-level median granule areas;$A_R$ =median of these nucleus-level values per animal.* | Median P granule area (px²) |
| **PGL-1 eccentricity** | PGL-1 | $$\widetilde{e_{R,k}} = \text{median}\left(e_{1,k}^R, \dots, e_{n_k^R,k}^R\right)$$ <br> $$E_R = \text{median}\left(\widetilde{e_{R,k}} : k \in V_R^e\right)$$ <br> $e_{i,k}^R = \sqrt{1-\left(b_{i,k}/a_{i,k}\right)^2}$ *where a and b are the semi−major and semi−minor axes of the fitted ellipse for granule (i) in nucleus k, respectively. Eccentricity is dimensionless: 0 indicates a circle, and values approaching 1 indicate increasing elongation; $n_k^R$ =number of PGL−1 granules in nucleus k;$V_R^e$ = set of nuclei with finite nucleus-level median eccentricities.* | Median shape elongation of P granules; higher values indicate greater elongation. |
| **MUT-16 foci count** | MUT-16 | $$\overline{N_G} = \frac{1}{K}\sum_{k=1}^{K} n_k^G$$ <br> $n_k^G$= *MUT-16 granule counts in nucleus k; K= total nuclei imaged per worm* | Mean Mutator focus count per nucleus |
| **MUT-16 focus area** | MUT-16 | $$\widetilde{a_{G,k}} = \text{median}\left(a_{1,k}^G, \dots, a_{n_k^G,k}^G\right), \quad n_k^G > 0$$ <br> $$A_G = \text{median}\left(\widetilde{a_{G,k}} : k \in V_G\right)$$ <br> *$a_{i,k}^G$= area (px²) of MUT-16 focus (i) in nucleus k;* <br> *$n_k^G$= number of MUT-16 foci in nucleus k; $\widetilde{a_{G,k}}$= nucleus-level median focus area;$V_G$ =set of nuclei with finite nucleus−level median focus areas;$A_G$ =median of these values per animal. Area values for nuclei with no detected MUT-16 foci are treated as missing and excluded from the animal-level area median. If no finite values remain, the animal-level area is missing.* | Median Mutator focus area (px²) |
| **MUT-16 eccentricity** | MUT-16 | $$\widetilde{e_{G,k}} = \text{median}\left(e_{1,k}^G, \dots, e_{n_k^G,k}^G\right), \quad n_k^G > 0$$ <br> $$E_G = \text{median}\left(\widetilde{e_{G,k}} : k \in V_G^e\right)$$ <br> $e_{i,k}^G = \sqrt{1-\left(b_{i,k}/a_{i,k}\right)^2}$, *where a and b are the semi-major and semi-minor axes of the fitted ellipse for MUT-16 focus i in nucleus k, respectively. Eccentricity is dimensionless: 0 indicates a circle, and values approaching 1 indicate increasing elongation. $n_k^G$= number of MUT-16 foci in nucleus k;$V_G^e$ =set of nuclei with finite nucleus−level median eccentricities. Eccentricity values for nuclei with no detected MUT-16 foci are treated as missing and excluded before animal-level aggregation. If no finite values remain, the animal-level value is missing.* | Median shape elongation of Mutator foci; higher values indicate greater elongation. |
| **PGL-1 NN-spacing** | PGL-1 | $$d_{i,k}^R = \min_{j \neq i} \lvert p_{i,k}^R - p_{j,k}^R \rvert_2$$ <br> $$C_{R,k} = \frac{\text{SD}\left(d_{1,k}^R, \dots, d_{n_k^R,k}^R\right)}{\text{mean}\left(d_{1,k}^R, \dots, d_{n_k^R,k}^R\right)}$$ <br> $$C_R = \text{median}\left(C_{R,k} : k \in V_R^{NN}\right)$$ <br> *$p_{i,k}^R$= peak−intensity position (x, y) of PGL−1 granule i in nucleus k; $d_{i,k}^R$= Euclidean distance from granule i to its nearest neighbor within nucleus k; $n_k^R$ = number of PGL-1 granules in nucleus k;$C_{R,k}$= nucleus−level nearest−neighbor spacing CV, calculated as sample SD divided by mean distance;$V_R^{NN}$ = set of nuclei with finite nucleus−level spacing CVs; $C_R$= median of these values per animal. Values for nuclei with fewer than four PGL-1 granules or a mean nearest-neighbor distance of zero are treated as missing and excluded from the animal-level median. If no finite values remain, the animal-level CV is missing.* | Nearest-neighbor spacing regularity of P granules; higher values indicate more variable spacing. Eglen et al. (2005) |
| **MUT-16 NN-spacing** | MUT-16 | $$d_{i,k}^G = \min_{j \neq i} \lvert p_{i,k}^G - p_{j,k}^G \rvert_2$$ <br> $$C_{G,k} = \frac{\text{SD}\left(d_{1,k}^G, \dots, d_{n_k^G,k}^G\right)}{\text{mean}\left(d_{1,k}^G, \dots, d_{n_k^G,k}^G\right)}$$ <br> $$C_G = \text{median}\left(C_{G,k} : k \in V_G^{NN}\right)$$ <br> *$p_{i,k}^G$ =peak−intensity position (x, y) of MUT−16 focus i in nucleus k; $d_{i,k}^G$=Euclidean distance from focus i to its nearest neighbor within nucleus k; $n_k^G$=number of MUT-16 foci in nucleus k; $C_{G,k}$=nucleus-level nearest-neighbor spacing CV, calculated as sample SD divided by mean distance; $V_G^{NN}$=set of nuclei with finite nucleus-level spacing CVs; $C_G$= median of these values per animal. Values for nuclei with fewer than four MUT-16 foci or a mean nearest-neighbor distance of zero are treated as missing and excluded from the animal-level median. If no finite values remain, the animal-level CV is missing.* | Nearest-neighbor spacing regularity of Mutator foci; higher values indicate more variable spacing. |

| | | | |
|---|---|---|---|
| **PGL-1 radial dispersion CV** | PGL-1 | $r^R_{i,k} = \lvert p^R_{i,k} - c_{img,k} \rvert_2$<br>$D_{R,k} = \frac{\mathrm{SD}\left(r^R_{1,k}, \ldots, r^R_{n^R_k,k}\right)}{\mathrm{mean}\left(r^R_{1,k}, \ldots, r^R_{n^R_k,k}\right)}$<br>$D_R = \mathrm{median}\left(D_{R,k}: k \in V^{rad}_R\right)$<br>*$p^R_{i,k}$ = peak−intensity position (x, y) of PGL−1 granule i in nucleus k; $c_{img,k}$ = image center, defined as (number of columns/2, number of rows/2); $r^R_{i,k}$ = Euclidean distance from granule i's peak−intensity position to the image center; $n^R_k$ = number of PGL−1 granules in nucleus k; $D_{R,k}$ = nucleus−level radial−distance CV, calculated as sample SD divided by mean radial distance; $V^{rad}_R$ = set of nuclei with finite radial−distance CVs; $D_R$ = median of these values per animal. Values are treated as missing when fewer than three PGL-1 granules are detected or the mean radial distance is less than $10^{-6}$ pixels. Missing values are excluded from the animal-level median; if no finite values remain, the animal-level CV is missing. Images were cropped to center each nucleus, so the image center was used as an approximation of the nuclear center.* | Relative variability in P granule radial distances from the nuclear centroid; higher values indicate greater relative variability in radial distances. |
| **MUT-16 → PGL-1 net association** | MUT-16 → PGL-1 | $Q_{G\to R,k} = 100\frac{n^{G,touch}_k}{n^G_k} - 100\frac{\lvert \mathrm{dil}(M_{R,k}, 7) \cap T_k \rvert}{\lvert T_k \rvert}$<br>$Q_{G\to R} = \mathrm{median}\left(Q_{G\to R,k}: k \in V_{G\to R}\right)$<br>*$n^{G,touch}_k$ = number of MUT-16 foci whose peak-intensity positions lie within the dilated PGL-1 mask in nucleus k; $n^G_k$ = total number of MUT−16 foci in nucleus k; $M_{R,k}$ = binary mask of segmented PGL−1 granules in nucleus k; $\mathrm{dil}(M_{R,k}, 7)$ = PGL-1 mask dilated using a disc-shaped structuring element of diameter 7 pixels; $T_k$ = foreground mask formed from pixels exceeding the corresponding image mean plus 0.5 SD in either channel, followed by dilation using a disc-shaped structuring element of diameter 5 pixels; $\lvert\cdot\rvert$ = number of pixels in a mask; $Q_{G\to R,k}$ = nucleus-level MUT-16 → PGL-1 net association, calculated as the percentage of MUT-16 peaks within the dilated PGL-1 mask minus the percentage of the foreground mask covered by that mask; $V_{G\to R}$ = set of nuclei with finite nucleus−level net−association values; $Q_{G\to R}$ = median of these values per animal. Net association is expressed in percentage points; positive values indicate association above the mask-based chance estimate. Values for nuclei with no detected MUT-16 foci are treated as missing and excluded from this feature's animal-level median. If no finite values remain, the animal-level value is missing. All quantities refer to one animal.* | Net % of Mut-16 foci proximal to P granules above tissue-level chance. Bolte & Cordelières (2006), Costes et al. (2004) |
| **PGL-1 → MUT-16 net association (norm.)** | PGL-1 → MUT-16 | $Q_{R\to G,k} = \frac{1}{n^G_k}\left[100\frac{n^{R,touch}_k}{n^R_k} - 100\frac{\lvert \mathrm{dil}(M_{G,k}, 7) \cap T_k \rvert}{\max(\lvert T_k \rvert, 1)}\right]$<br>$Q_{R\to G} = \mathrm{median}\left(Q_{R\to G,k}: k \in V_{R\to G}\right)$<br>*$n^{R,touch}_k$ = number of PGL-1 granules whose peak-intensity positions lie within the dilated MUT-16 mask in nucleus k; $n^R_k$ = number of PGL−1 granules in nucleus k; $n^G_k$ = number of MUT−16 foci in nucleus k; $M_{G,k}$ = binary mask of segmented MUT−16 foci; $\mathrm{dil}(M_{G,k}, 7)$ = MUT−16 mask dilated using a disc−shaped structuring element of diameter 7 pixels; $T_k$ = foreground mask formed from pixels exceeding the corresponding image mean plus 0.5 SD in either channel, followed by dilation using a disc-shaped structuring element of diameter 5 pixels; $\lvert\cdot\rvert$ = number of pixels in a mask; $Q_{R\to G,k}$ = nucleus−level net association divided by MUT−16 focus count; $V_{R\to G}$ = set of nuclei with finite normalized net−association values; $Q_{R\to G}$ = median of these values per animal. Values are treated as missing when either channel has no detected objects. The animal-level median uses finite values only and is missing if none remain. All quantities refer to one animal. The PGL-1 → MUT-16 index was divided by MUT-16 focus count to express net association relative to focus abundance, because MUT-16 focus number varies between nuclei, whereas PGL-1 granule number—the target abundance in the reverse index—is comparatively stable.* | As above for P granules near Mutator foci, divided by $n^G_k$ to normalize for focus abundance |
| **PGL-1 / MUT-16 size ratio** | Both | $S_k = \frac{\widetilde{a_{R,k}}}{\widetilde{a_{G,k}}}$<br>$S = \mathrm{median}(S_k: k \in V_S)$<br>*$\widetilde{a_{R,k}}$ = median PGL-1 granule area in nucleus k ($px^2$); $\widetilde{a_{G,k}}$ = median MUT−16 focus area in nucleus k ($px^2$); $S_k$ = ratio of the nucleus-level median PGL-1 granule area to the nucleus-level median MUT-16 focus area; $V_S$ = set of nuclei with finite area ratios; S = median of these nucleus−level ratios per animal. Ratios are treated as missing when either channel has no detected objects, either median area is nonfinite, or the median MUT-16 focus area is zero. The animal-level median uses finite ratios only and is missing if none remain. All quantities refer to one animal. The ratio is dimensionless.* | Relative granule size; >1 means P granules larger than Mutator foci. |

**Age-associated changes in germ-granule architecture**
Age effects on the twelve clock features were tested in Batch 1 and Batch 2 AL worms using linear mixed-effects models with a random intercept per animal, because nuclei were nested within animals (feature ~ age + (1 | animal); lme4 (v2.0-1), lmerTest v3.2.1). Features were z-scored, so slopes are in SD per day. Non-finite values were treated as missing. MUT-16 focus area, MUT-16 eccentricity, and MUT-16 → PGL-1 net association are undefined in images without MUT-16 foci, so they were set to missing there. A sensitivity analysis kept these images as zeros. Models were fitted to each batch and to the combined data (batch as a fixed effect), for features with at least three animals on every day. Slopes are reported with Wald 95% CIs and Satterthwaite t-test p-values (combined model, Table S1; Batch 1, Table S2; Batch 2, Table S3). The overall effect of day was also tested with age as a categorical factor (type III F-test), and batch-by-age interaction by likelihood-ratio test (ML fits; Table S4). P values were adjusted across features within each model by Benjamini–Hochberg (BH), with BH-adjusted $P < 0.05$ considered significant. The combined model defined the age-associated features.

**GG-Clock construction and validation**
Chronological age was modeled from the twelve animal-level features by elastic net regression (Zou & Hastie, 2005), with each animal as the unit of analysis, following the penalized-regression approach widely used for aging clocks (Horvath, 2013). Chronological age (days) was regressed on the twelve features with glmnet (v5.0; Gaussian; $\alpha = 0.5$; internal standardization). $\lambda$ was chosen from 100 log-spaced values ($10^2$ to $10^{-5}$) by lowest mean squared error in 10-fold cross-validation with random folds (seed 19 for the final clock; no one-standard-error rule). Missing values were imputed with training medians, and features that were all-missing or constant in training were given weight 0. The final clock was fitted once on all ad libitum (AL) animals from Batch 1 (discovery cohort). Feature importance was the coefficient × training SD, and each trait's share was its absolute standardized coefficient divided by the sum over all traits (Table S7).

Performance was assessed by leave-one-worm-out cross-validation (LOO-CV) (Hastie et al., 2009; Meyer & Schumacher, 2021). In each fold, imputation, $\lambda$ tuning, and fitting were repeated on the remaining animals, so each animal was predicted by a model fitted without it. Predictions were left unclipped. Performance was summarized as MAE, RMSE, and bias (predicted minus chronological age), overall and per day, with overall Spearman $\rho$ and Pearson's $r^2$ (squared Pearson correlation) (Table S5). Selection frequency was the percentage of LOO fits with a non-zero coefficient (Table S7). To quantify variability from random fold assignment, the full LOO procedure was repeated with 30 seeds (1 to 30), and the mean, SD, and range of $\rho$, $r^2$, and MAE are reported (Table S6). For the extended clock, which included day 6 animals, the model was constructed, cross-validated, and evaluated by the same procedure as the primary clock (days 1–5) (Table S26).

To test whether the GG-Clock responds to a non-genetic longevity intervention, animals subjected to intermittent fasting (IF) were segmented using the Batch 1 AL image-noise reference and projected onto the frozen clock using the training-derived imputation medians and scaling. IF animals were not used for training in either batch. The final Batch 1 clock was applied unchanged to Batch 2 AL and IF animals and was not refit or tuned on Batch 2. The two batches were analyzed separately, without pooling. In Batch 1, same-day AL controls were represented by their LOO predictions. External performance in Batch 2 AL animals was summarized as Spearman $\rho$, Pearson's $r^2$, MAE, and RMSE (Table S12; per-animal predictions in Table S17). IF animals were compared with concurrently imaged AL controls from the same replicate. The effect of IF on prediction error (predicted minus chronological age) was estimated in each batch by linear regression (Batch 1, Table S8; Batch 2, Table S13). The IF − AL difference is reported with a t-based 95% CI and a one-sided P-value (IF younger). Per-day mean errors are shown descriptively. Feature contributions to the IF shift were calculated with fixed AL clock weights as weight × the difference in mean imputed values (IF minus AL). These sum to the difference in mean predicted age. They were calculated at day 4 in both Batches 1 and 2 (Tables S9 and S16). Uncertainty was estimated by an animal-level bootstrap (5,000 resamples; seed 19; AL and IF resampled independently) with 95% percentile CIs. Clock-training uncertainty was not propagated.

**Brood size assay**
Worms were synchronized by bleaching. At mid-L4, 20 worms were singled onto individual 35-mm NGM plates seeded with OP50-1 and transferred to fresh plates daily until reproduction ceased. Plates were kept at 20 °C for 3 d, until progeny reached the first day of adulthood, and progeny were then counted per worm per day (Days 1 to 6). Five animals were excluded because of bagging, leaving 15 animals with complete records. This cohort was a parallel subset of the same synchronized population used for imaging, and the clock data came from separate imaged animals (Batch 1 leave-one-worm-out predictions, Days 1 to 6).

Daily progeny are summarized as mean, SD, median, and SEM per day for all worms with a record on each day (n = 16 on Days 1 to 3 and n = 15 on Days 4 to 6; Table S18). Cumulative progeny were calculated per worm for the 15 worms with complete Day 1 to 6 records (Table S20), and lifetime brood was the sum of daily progeny (Table S21). Cumulative output was also expressed as a percentage of each worm's own lifetime brood, excluding worms with zero brood from this calculation. The plateau was defined as the first day on which the mean of the per-worm cumulative percentages reached at least 95% of the lifetime brood. Changes in daily progeny with age were tested on Days 1 to 5 using a negative binomial generalized linear mixed model (glmmTMB 1.1.15.2; nbinom2) with day (factor) as a fixed effect and worm as a random intercept. Day 6 was excluded from the model because all counts were zero. The effect of day was assessed by a likelihood-ratio test against an intercept-only model with the same random effect. Each day was compared with Day 1 by Dunnett-adjusted contrasts of estimated marginal means (emmeans 2.0.3), reported as response ratios with 95% CIs (likelihood-ratio test, contrasts, and estimated means, Table S19). The clock and progeny data were compared descriptively by day, at cohort level, without a correlation test (Table S22). Changes in mean predicted age between days were estimated as later-day minus earlier-day differences, with 95% percentile bootstrap CIs (5,000 resamples of animals within each day; seed 19) and one-sided Mann-Whitney U tests (later day > earlier day; unadjusted P values) (Table S23).

**GGProfiler software**
GGProfiler (v1.0) is an open-source, browser-based tool written in R with the Shiny (v1.14.0) framework (EBImage for image processing; ggplot2 v4.0.3 and DT v0.34.0 for plots and tables). It implements the segmentation and feature-extraction pipeline described above (see Image processing and segmentation) to profile germ-granule morphology and spatial organization in individual germline nuclei.

Setup supports one to four user-defined channels, each with a marker name, display color, segmentation method (top-hat or DoG, as described above), and RGB-plane selector (ignored for single-plane grayscale files). Analysis steps accept single 2D images, so z-stacks must be converted beforehand. An optional Crop nuclei module reads TIFF, PNG, or JPEG files. The user manually draws a rectangle around each nucleus, and the same rectangle is applied to every channel image of the field. The module writes 16-bit TIFFs and a crop log with the coordinates. In the Conditions tab, channel files are paired within each condition by the trailing numeric ID in the file name. A pairing manifest labels each file as ready, excluded (missing or duplicate channel file), or invalid (no numeric ID). An interactive QC module previews one sample at a time, showing a display-normalized composite, segmentation outlines, and an annotated overlay with a context plot and a table of preview features.

From the retained objects, GGProfiler computes five per-channel measurements: object count (_n), median object area (_size), median eccentricity (_elong), nearest-neighbor distance CV between intensity peaks (_ring; at least four objects), and CV of peak distances from the image center (_radcv; at least three objects). For every channel pair (A, B), it also computes chance-corrected B-to-A net proximity (<B>_near_<A>_net, in percentage points), A-to-B net proximity (chance-corrected, then divided by the B object count; <A>_near_<B>_net_norm, so it is not on the same scale as the B-to-A value), and the ratio of median A to median B object area (_sizeratio). Proximity metrics use masks dilated by 7 px, with chance expectation estimated within foreground (pixels above mean + 0.5 SD in any channel). GGProfiler also offers an optional, direct measure of object overlap (Khushi et al., 2017), which is calculated on undilated masks and was not used in the clock analyses of this study. It gives a directly interpretable readout of physical contact between granules of different channels. An A–B object pair is counted as overlapping when the two objects share at least a user-set minimum number of pixels (default 3). GGProfiler reports the number of overlapping pairs (_segcoloc_pairs_n) and the percentage of objects in each channel that overlap at least one object of the other channel (_segcoloc_b_to_a_pct, _segcoloc_a_to_b_pct). Each channel pair is analyzed separately, and values are not pooled. Spatial measurements are in pixels.

The Extract tab processes all paired samples and exports per-image features, condition summaries (counts averaged, other features by median), per-object measurements, a processing log, and an RDS snapshot of results, labels, settings, calibration, and session information.

The Compare tab tests per-image features between two selected conditions, with the image as the unit of observation. Granule-level rows and condition summaries are not tested. For each feature, non-finite values are excluded, and a test is run only when each condition has at least three finite observations. The user selects the statistical test: either a two-sided Wilcoxon rank-sum test (normal approximation, exact = FALSE) or a two-sided Welch t-test (unequal variances). Effect sizes are reported as the difference in group means and medians (condition B minus condition A) in the feature's original units. P-values are adjusted across all features in the selected condition pair by the Benjamini-Hochberg procedure, and a feature is called significant when its adjusted P-value is below a user-defined threshold α (default 0.05). The tests assume independent observations and do not account for multiple images from the same animal or sample. In this study, the Wilcoxon rank-sum test was used with α = 0.05 to compare control images with heat-shocked images (Table S24) and with starved images (Table S25). Results are exported as CSV.

The Visualize tab produces distribution plots (box, violin, or mean ± SE) and PNG or PDF images of raw composites, segmentations, and annotated overlays. The Figure export tab produces display-only PDFs of the raw channels with adjustable limits, brightness, contrast, and colors, without altering the stored data. Further details are given in the GGProfiler manual. GGProfiler can be accessed at https://github.com/Purbashadg/Germ-Granule-Remodeling.

**Statistical analysis**

Statistical analyses were performed in R (v4.5.0), and figures were generated with ggplot2 (Wickham, 2016). Statistical methods are described in their respective sections above.

**Use of AI declaration**

During the preparation of this work, the authors used ChatGPT (GPT-6 Astra; OpenAI) to improve the flow of the manuscript, to condense text, and to troubleshoot code.
