## Supplementary material for "Germ Granule Remodeling Reveals Germline Age in *Caenorhabditis elegans*": GGProfiler v1.0 user manual

### GG Profiler

GERM GRANULE PROFILER

*Multi-channel germ granule analysis and spatial profiling*

**User Manual and Technical Reference**

**Software version:** GGProfiler v1.0
**Source code and downloads:** https://github.com/Purbashadg/Germ-Granule-Remodeling
https://www.thesglab.com/resources
**Implementation:** R / Shiny; image analysis with EBImage
**Developed by:** The Silva-Garcia Lab

This manual describes the workflow, the algorithms, and every exported measurement of GGProfiler, so that analyses can be reproduced and reported accurately in Methods sections.

**Content**

1. Overview and intended use
2. Installation and launching
3. Image requirements and file naming
4. Workflow at a glance
5. Setup tab
6. Crop nuclei tab
7. Conditions tab
8. QC tab: calibration, segmentation and overlays
9. Extract tab and output files
10. Compare tab (statistics)
11. Visualize tab
12. Figure export tab
13. Algorithms in detail
14. Measurement reference
15. Parameter reference and defaults
16. Reproducibility and reporting guidance
17. Limitations and interpretation cautions
18. Troubleshooting
19. References

#### 1. Overview and intended use

GGProfiler is an open-source (source code: https://github.com/Purbashadg/Germ-Granule-Remodeling), interactive application that automates the segmentation and quantification of germ granule compartments (for example, PGL-1:mCardinal P granules and MUT-16:GFP Mutator foci) in single-nucleus fluorescence images. For every nucleus, it detects granules in each marker channel, then measures granule number, size, shape, spatial regularity, distance from the image center, and between-channel spatial relationships. It then supports statistical comparison of experimental conditions and export of publication-style figures.

The application performs the following steps: (i) cropping of single nuclei from field images, (ii) pairing of per-marker images by nucleus ID, (iii) quality-controlled segmentation with live overlays, (iv) batch feature extraction, (v) two-condition statistical comparison, and (vi) figure and data export. All spatial measurements are expressed in pixels.

**Instructions panel.** Each tab of the application contains an Instructions panel that explains the controls and recommended steps for that tab. Read the Instructions of each tab before using it and use them together with this manual.

#### 2. Installation and launching

##### 2.1 Requirements

| Package | Source | Status | Use |
|---|---|---|---|
| shiny | CRAN | Required | Application framework |
| ggplot2 | CRAN | Required | Distribution plots, crop preview |
| DT | CRAN | Required | Interactive tables |
| EBImage | Bioconductor | Required | Image reading, morphology, labeling, shape features |
| zip | CRAN | Optional | Safer ZIP export of cropped images (falls back to utils:zip) |
| ragg | CRAN | Optional | Higher-quality PNG export (falls back to base png) |

On start-up the script checks for missing packages. In an interactive session, it offers to install them (CRAN and Bioconductor via BiocManager); otherwise, it stops with explicit installation commands. To reproduce the exact package versions used in the study, open GGProfiler.Rproj in RStudio and run renv::restore() (versions are recorded in renv.lock).

##### 2.2 Launching

1. Download the repository from GitHub (https://github.com/Purbashadg/Germ-Granule-Remodeling). The application is in the GGProfiler folder, which contains the script 06_GGProfiler.R, the www subfolder (logos), and the renv.lock file listing the package versions used.
2. Optional: place logo files (image.png, Lab Logo full.png) in a www subfolder. They affect only the header; missing logos do not affect analysis.
3. In R or RStudio run shiny::runApp("path/to/GGProfiler/06_GGProfiler.R")(replace the path with the location of your folder), or open 06_GGProfiler.R in RStudio and click Run App.

On first launch, GGProfiler checks for required packages (shiny, ggplot2, DT, EBImage) and offers to install any that are missing. Type **y** when prompted. This needs an internet connection and R ≥ 4.1.
The upload limit is 2 GiB per request. Uploaded files are copied to a per-session temporary folder that is deleted when the session ends. Folder-based saving of crops (Section 6) works only when the application runs on the user's own computer.

#### 3. Image requirements and file naming

- **Format:** TIFF, PNG or JPEG. Single 2D grayscale images (recommended) or RGB images. Z-stacks and time series are not accepted by the analysis tabs; generate a 2D projection or select a plane first. Images must be at least 7 x 7 pixels and contain only finite values.
- **Matching crops:** all channels of one nucleus must have identical pixel dimensions and be aligned. Crop each nucleus carefully along its periphery (Section 6).
- **Grayscale vs. RGB:** for grayscale files the single plane is used and the RGB source setting is ignored. For RGB files, the selected R, G or B plane is analyzed. Composite images can be used only when each marker occupies a distinct RGB plane; pseudocolored composites that mix markers across planes should not be used.
- **File naming:** the nucleus ID is the number at the very end of the file name, before the extension (for example PGL1_1.tif and MUT16_1.tif are nucleus 1). Pairing is by this number, not by upload order. Leading zeros are ignored (001 equals 1). Use exactly one file per marker per nucleus and a unique ID per nucleus within each condition (including across animals and fields). IDs may be reused between conditions.

#### 4. Workflow at a glance

| Step | Tab | Action | Result |
|---|---|---|---|
| 1 | Setup | Define 1-4 channels: name, RGB source, display color, segmentation method; click Apply setup | Channel configuration |
| 2 | Crop nuclei (optional) | Draw a rectangle around each nucleus; save crops | Matched per-marker TIFFs, crop log |
| 3 | Conditions | Add conditions; upload images per marker | Pairing manifest (ready / excluded / invalid) |
| 4 | QC | Calibrate noise (DoG channels); tune parameters; preview segmentation on representative nuclei | Validated parameter set |
| 5 | Extract | Run extraction on all paired nuclei | Per-image, summary, object and log tables |
| 6 | Compare | Choose a condition pair, test and correction | Per-feature statistics |

| | | | |
|---|---|---|---|
| 7 | Visualize / Figure export | Plot distributions; export overlays and display figures | PNG/PDF figures |

#### 5. Setup tab

**Instructions:** See the Instructions panel within the Setup tab for step-by-step guidance on its controls. Choose the number of channels (1-4), then configure a card for each channel:

| Setting | Meaning |
|---|---|
| Marker name | Label used in upload fields and in exported column names. Names must be non-empty and unique, including after R name sanitization (make.names), so "PGL-1" becomes PGL.1 in column names. |
| RGB source | Plane (R, G or B) read from RGB images. Ignored for grayscale files. |
| Display color | Color used in overlays only. It never changes measured intensities. |
| Segmentation | Top-hat (diffuse or larger objects) or DoG (punctate objects). Details in Section 13. |

The default setup is two channels: channel 1 "PGL-1" (red, top-hat) and channel 2 "MUT-16" (green, DoG). Click Apply setup to commit the choices. Changing the setup after conditions exist requires removing all conditions first; editing the cards without applying marks the setup as unapplied and blocks analysis.
**Channel order matters.** Pairwise measurements are directional. The first channel is "A" and the second is "B" in every pair (Section 14). With the default setup A = PGL-1 and B = MUT-16.

#### 6. Crop nuclei tab

**Instructions:** See the Instructions panel within the Crop nuclei tab for step-by-step guidance on its controls. This optional tab cuts single nuclei out of a larger field and creates matched per-marker images for the Conditions tab. Cropping uses raw pixel values; the on-screen contrast stretch (0.1th to 99.9th percentile) is for viewing only.

**Crop carefully along the nuclear periphery.** The crop is the only definition of the nucleus in GGProfiler. Draw each rectangle tightly and very carefully along the periphery of the nucleus, so that the nucleus fills the crop and its center coincides with the image center. Loose or off-center crops add background, can include neighboring nuclei, and directly change the radial measurement (_radcv), nearest-neighbor spacing and the chance-corrected proximity values. Apply the same cropping criterion to every nucleus, condition and cohort, and report it in Methods.

1. Upload all images of one field together (one grayscale file per marker, or one merged RGB image). Images whose dimensions differ from the reference image are skipped and reported.
2. Choose the reference image used for drawing. For multi-plane TIFFs a Z-plane slider selects the displayed plane (the crop applies to all planes).
3. Zoom (optional): drag a box, click Zoom to dragged box; use Zoom out or Reset zoom. Zoom changes the view only.
4. Draw a rectangle around one nucleus and click Apply rectangle (minimum 7 x 7 pixels).
5. Save crop with a unique nucleus number. The same rectangle is applied to every uploaded image. Saved rectangles are outlined in cyan; Remove last and Clear all undo saves.

#### Outputs

- File names are <source file stem>_<nucleus number>.tif, saved as 16-bit TIFF without intensity rescaling.
- Save to folder writes TIFFs and a time-stamped crop_log_*.csv directly to a chosen folder (local use only). Download ZIP bundles TIFFs and crop_log.csv. Existing files are skipped unless Overwrite is ticked. By default all planes of a multi-plane TIFF are saved; tick Save only the displayed plane for 2D output.
- The crop log records source file, output file, rectangle coordinates (EBImage convention: x = column, y = row, 1-based, y = 1 is the top row), planes saved, status and application version. Keep it with the data and report the cropping criterion in Methods.

Cropping does not align images and does not detect nuclei automatically.

#### 7. Conditions tab

**Instructions:** See the Instructions panel within the Conditions tab for step-by-step guidance on its controls.

1. Enter a condition name (for example "AL day 4" or "IF day 4") and click Add condition (maximum 20; names must be unique).
2. In the condition card, upload the images for each marker using that marker's field. Select all files for the marker at once. Uploading again replaces the previous selection for that marker and condition; it does not append.
3. For merged RGB images, upload the same complete batch under every marker, with filenames unchanged.
4. Inspect the pairing manifest.

| Status | Description |
|---|---|
| ready | Exactly one file exists for every configured marker for that nucleus ID. |
| excluded | A marker file is missing, or a numeric ID is duplicated within a marker. See the detail column and correct the upload. |
| invalid | The file name does not end in a numeric ID. Rename and upload again. |

"Ready" confirms file-name pairing only. It does not check alignment, image quality or the chosen RGB plane; dimension checks occur at analysis time. The manifest can be downloaded as CSV. Changing uploads or conditions makes previous calibration and extraction results stale.

#### 8. QC tab: calibration, segmentation and overlays

**Instructions:** See the Instructions panel within the QC tab for step-by-step guidance on its controls.
QC previews one sample at a time and never replaces full extraction. Inspect representative nuclei from every condition before extracting.

##### 8.1 Controls

| Control | Function |
|---|---|
| Sample | Selects a paired nucleus (condition / nucleus ID). |
| Use legacy batch-noise scaling | When enabled (default), the local-contrast cutoff of every DoG channel is scaled by the sample's background noise relative to the dataset reference. It does not normalize intensities and does not affect top-hat |

| | |
|---|---|
| | channels. When enabled, calibration is mandatory for preview and extraction. |
| Calibrate current dataset | Estimates a noise reference for each DoG channel from all complete paired samples (Section 13.4). Any change of channel settings, parameters or uploads makes calibration stale. |
| Preview segmentation | Runs segmentation and measurement on the selected sample with the current settings. |
| Overlay | Selects the annotation in the third panel and its context plot (below). |
| Distance-overlay channel | Channel used for nearest-neighbor links and radial spokes. |
| Channel pair A / B | Channels used for proximity and segmented-overlap displays. Order defines direction. |
| Minimum shared pixels per object pair | Threshold for segmented-object overlap (default 3 px). Changing it requires a new preview and extraction, but not recalibration. |

#### 8.2 Display panels

- **Raw panel:** display-normalized composite of all channels (each channel scaled min-max, for viewing only).
- **Segmentation panel:** composite with outlines of retained objects.
- **Annotated panel and context plot:** the selected overlay and its supporting summary.
- **Preview features table:** all feature values for the previewed sample. NA means undefined, not zero.

#### 8.3 Overlays

| Overlay | Shows |
|---|---|
| Peak positions | Brightest pixel inside each object (not necessarily the centroid). Context plot: object counts per channel. |
| Nearest-neighbor links | Lines joining each peak to its nearest same-channel peak (peak-to-peak distance, not edge-to-edge). Needs at least two objects. |
| Radial spokes: image center | Lines from the image center to each peak. The reference is the center of the image, not a segmented nuclear boundary or nuclear center; crop nuclei so that the nucleus is centerd. |
| B peaks near A objects | B peaks tested against the dilated A mask. Circles = within the mask; crosses = outside. Reports the chance-corrected net value. |
| A peaks near B objects | A peaks tested against the dilated B mask. The reported value is chance-corrected and divided by the number of B objects. |
| Segmented-object overlap | White pixels mark shared pixels of A-B object pairs that meet the minimum-pixel threshold (masks not dilated). |

##### 8.4 Recommended QC sequence

1. Confirm channel setup and sample pairing.
2. Choose a sample and review parameters (Section 15).
3. Calibrate if batch-noise scaling is enabled and any channel uses DoG.
4. Click Preview segmentation; compare raw and segmentation panels for missed granules, background objects, merged or fragmented objects.
5. Inspect overlays and numerical features; repeat on samples from each condition (and from young and old animals when relevant).
6. Freeze parameters before extraction. Use the same parameters for every condition in a comparison.

#### 9. Extract tab and output files

**Instructions:** See the Instructions panel within the Extract tab for step-by-step guidance on its controls.
Click Run extraction to process every ready sample with the current settings. Failed images are logged and skipped; the summary reports processed, successful, failed and zero-object counts. Results are valid only for the settings at extraction time; if settings change, the tab reports that previous results are stale.

| Output | Contents |
|---|---|
| Per-image CSV | One row per nucleus: condition, image name and all features (Section 14). |
| Condition summary CSV | Per condition: number of images; count features (_n) are summarized as the mean, all other features as the median, using finite values only. |
| Object measurements CSV | One row per detected granule: channel, object ID, area (px), peak and centroid coordinates, eccentricity, mean and peak intensity, local background median and MAD, local-contrast z, nearest-neighbor distance and radial distance (px). |
| Processing log CSV | Per image: status (ok, ok_zero_objects, failed), error and warnings. |
| Analysis snapshot RDS | Results, labels, settings, calibration, manifest, statistics, session information and notes. It is a record, not a project importer, and does not embed the original images. |

Archive the per-image CSV, object CSV, processing log, crop log, manifest and snapshot with each dataset. Together they document every parameter and exclusion.

#### 10. Compare tab (statistics)

**Instructions:** See the Instructions panel within the Compare tab for step-by-step guidance on its controls.
The Compare tab tests each feature between two conditions using per-image (per-nucleus) values. Differences are always B minus A.

| Option | Behavior |
|---|---|
| Condition pair | Any two conditions with extracted data (A = first listed, B = second). |

| | |
|---|---|
| Test | Wilcoxon rank-sum: two-sided, normal approximation (exact = FALSE). Welch t-test: two-sided, unequal variances. |
| BH correction | Benjamini-Hochberg across all features of the selected pair (default on). If off, adj_p equals the raw p-value and the correction column reads "none". |
| Alpha | A feature is "significant" when finite adj_p < alpha (default 0.05). |

For each feature, non-finite values are removed and at least three finite observations per condition are required, otherwise an error message is recorded and no p-value is produced. The results table reports test, n per group, means, medians, delta_mean, delta_median, raw p-value, adjusted p-value, correction, alpha, significance flag, and any error or warning. The difference plot shows median differences (unstandardized, in feature-specific units) for significant features only; bar lengths from different features are not comparable.

**Statistical independence.** Both tests assume independent observations. Nuclei from the same animal are correlated, and the Compare tab does not model animals as a grouping factor. For publication, export the per-image table and analyze nuclei nested within animals (for example a linear mixed model with a random intercept for worm, or analysis of per-animal means), and report the unit of replication. Use the Compare tab for exploration and quality checks.

#### 11. Visualize tab

**Instructions:** See the Instructions panel within the Visualize tab for step-by-step guidance on its controls.

- **Distribution plot:** choose a feature and a plot type: box + points, violin + points (groups with at least two points), or mean + SE. If a current comparison exists, the plot caption shows the pair, test, correction and adjusted p-value. "Significant features only" filters the feature list.
- **Browser:** shows the raw, segmentation and annotated panels of any extracted image from stored results (no new segmentation).
- **Export:** PDF (Helvetica, vector, default) or PNG. Controls: width (mm), distribution-plot height (mm), PNG resolution (72-1200 dpi, default 600). Export figures use publication styling (Helvetica, 11 pt text, 10 pt ticks, 0.5 pt lines, 2.4 pt points). Raw and segmentation images can be exported as clean images (no title; one image pixel equals one output pixel in PNG) or as annotated overlays with title and caption.

#### 12. Figure export tab

**Instructions:** See the Instructions panel within the Figure export tab for step-by-step guidance on its controls.
This tab (added after the main tabs load) produces display-only figure panels from the original image channels. It runs no segmentation and changes no measurement; it needs only uploaded, paired samples.

- **Per channel:** show/hide, figure color, base display minimum and maximum, brightness (-100 to +100%) and contrast (0.1x to 5x). Display minimum maps to black, display maximum to full channel color; values outside are clipped for display. Brightness shifts and contrast narrows or widens the display window.
- Auto contrast sets limits to the 0.5th and 99.5th percentiles of the selected image; Reset uses the full intensity range. Both affect all channels and reset brightness and contrast.
- Limits and colors persist when switching samples. For comparable figures use identical limits for each marker across images acquired under the same settings. Applying Auto separately to each image can hide real intensity differences.

- A histogram of raw intensity with dashed lines at the display limits and a clipping table (percentage of pixels below minimum and above maximum) support honest reporting.
- **PDF export:** width in mm (10-500); height follows the image aspect ratio. The PDF embeds an RGB display raster at the native pixel dimensions without interpolation; it neither adds resolution nor preserves quantitative intensity precision. The effective resolution (ppi) is displayed.

State in figure legends that images were adjusted for display only and that quantification used unadjusted data.

### 13. Algorithms in detail

#### 13.1 Image reading

Images are read with EBImage. A 2D image is used as is. An RGB image yields the plane selected in Setup. Values are used as loaded (no rescaling) for segmentation and measurement.

#### 13.2 Top-hat segmentation

1. Gaussian blur with sigma = 1 pixel.
2. Background estimate by morphological opening with a disc brush of the user-set diameter (default 51 px; capped to fit the image and forced odd).
3. Response = max(blurred image - background, 0).
4. Candidate threshold = mean of positive response values + threshold multiplier x their standard deviation.
5. Binary mask is hole-filled, closed with a 3-px disc brush and connected components are labeled.
6. Objects are kept if minimum area ≤ area ≤ maximum area (pixels).

#### 13.3 DoG (difference-of-Gaussians) segmentation

1. Response = max(Gaussian(fine sigma) - Gaussian(coarse sigma), 0). The coarse sigma is capped at one sixth of the smaller image dimension and must exceed the fine sigma.
2. Candidates are obtained exactly as in steps 4-5 of top-hat segmentation.
3. An object is retained only if all of the following hold:
    - area between minimum and maximum;
    - mean raw intensity ≥ image mean + global floor multiplier x image SD;
    - eccentricity ≤ maximum eccentricity (EBImage moment eccentricity, 0-1);
    - local-contrast z ≥ local-contrast z minimum.

Local-contrast z is computed per object: the background is the ring of pixels within 4 px outside the object (dilation with a disc of diameter 9). z = (object mean intensity - ring median) / (1.4826 x ring MAD), with the MAD floored at 1e-9. Rings require at least five pixels. In the retained legacy behavior, neighboring objects are not excluded from the ring, so crowded regions can affect z.

#### 13.4 Batch-noise calibration and scaling

For each DoG channel and sample, the noise scale is the median ring MAD across candidate objects detected with a fixed multiplier of 2 (if none can be computed, a fallback of 1 is used with a warning). Calibration takes the median of the noise scales across all complete samples as that channel's reference. During preview or extraction with scaling enabled, the effective local-contrast cutoff for a sample is:

| effective z_min = entered z_min x (sample noise scale / reference noise scale) |
|---|

Thus noisier images receive a stricter cutoff. Top-hat channels are not scaled. Calibration is not an intensity-standardization procedure and does not guarantee comparable acquisition settings.

#### 13.5 Peaks and distances

- **Peak:** the pixel with maximum intensity within each object (first in index order if tied).

- **Nearest-neighbor distance:** Euclidean peak-to-peak distance to the closest peak of the same channel, computed row-wise.
- **Radial distance:** Euclidean distance from each peak to the image center ((width+1)/2, (height+1)/2).

##### 13.6 Proximity (dilation-based association)

1. Each channel's mask is dilated with a disc brush of diameter 7 px (capped to image size).
2. The tissue domain is the union over channels of pixels brighter than channel mean + 0.5 SD.
3. Chance for a channel = 100 x (dilated mask pixels inside the tissue domain) / (tissue domain pixels).
4. Gross association = percentage of the other channel's peaks lying inside the dilated mask.
5. Net = gross - chance, in percentage points.

Both channels must contain at least one object for pairwise values to be defined. Because the tissue domain is a union of all configured channels, adding or removing a channel can change chance-adjusted values.

##### 13.7 Segmented-object overlap

Inspired by MatCol (Khushi et al. 2017), this counts shared pixels between individual labeled objects of two channels, using the original masks without dilation or additional thresholding. A pair qualifies if it shares at least the minimum number of pixels (default 3); overlaps with different partners are never pooled. The pair count includes every qualifying pair; directional percentages count each source object once. If a source channel has no objects the percentage is NA; if source objects exist but none overlap it is 0. MatCol's background thresholding, Monte Carlo randomization and within-image significance testing are not reproduced. This is two-dimensional overlap and not evidence of molecular interaction.

#### 14. Measurement reference

Column names combine the marker name (sanitized, for example PGL.1) with a suffix. For pairs, the prefix is <A>_vs_<B>, with A and B in channel order.

##### 14.1 Per-channel features

| Suffix | Definition | Units / requirements |
|---|---|---|
| _n | Number of detected objects | Count; may be 0 |
| _size | Median object area | Pixels; NA if no objects |
| _elong | Median EBImage moment eccentricity (0 = round, 1 = line) | Unitless; NA if no objects |
| _ring | Coefficient of variation (SD/mean) of nearest-neighbor peak distances. A legacy name: it measures spacing regularity, not ring detection. Higher = more irregular spacing. | Unitless; needs ≥ 4 objects |
| _radcv | Coefficient of variation of peak distances from the image center | Unitless; needs ≥ 3 objects |

##### 14.2 Pair features (per ordered channel pair A, B)

| Suffix | Definition | Units / requirements |
|---|---|---|

| | | |
|---|---|---|
| <B>_near_<A>_net | B peaks within dilated A mask (gross %) minus A chance % | Percentage points; both channels need objects |
| <A>_near_<B>_net_norm | (A peaks within dilated B mask, gross % minus B chance %) divided by the number of B objects | Percentage points per B object; both channels need objects |
| _sizeratio | Median A object area divided by median B object area | Unitless; B size must be > 0 |
| _segcoloc_pairs_n | Number of qualifying A-B object pairs | Count |
| _segcoloc_b_to_a_pct | Percentage of B objects overlapping at least one A object | %; NA if B has no objects |
| _segcoloc_a_to_b_pct | Percentage of A objects overlapping at least one B object | %; NA if A has no objects |

**Linking to the manuscript feature set.** With the default setup (A = PGL-1, B = MUT-16), the per-channel suffixes and pair suffixes correspond to the granule traits used in the GG-Clock analysis scripts (counts, size, elongation, spacing regularity, radial dispersion, apposition and size ratio).

#### 14.3 Missing and zero values

NA (undefined) is not zero. Object counts of zero are valid values. Features that depend on objects (size, elongation, spacing, apposition) are NA when objects are absent or too few. Downstream analyses must state how NA values are handled; in the associated study, MUT-16-dependent features were masked (set to NA) when no MUT-16 foci were detected, before aggregation.

### 15. Parameter reference and defaults

| Parameter | Applies to | Default | Effect |
|---|---|---|---|
| Threshold multiplier | All | 1 (channels 1, 4); 2 (channels 2, 3) | Candidate threshold = mean positive response + multiplier x SD. Higher is stricter and can change merging and splitting. |
| Minimum area (px) | All | 10 (ch 1, 4); 20 (ch 2, 3) | Rejects smaller objects. |
| Maximum area (px) | All | 3000 (ch 1, 4); 1000 (ch 2, 3) | Rejects larger objects; does not split merged objects. |
| Top-hat brush diameter (px) | Top-hat | 51 | Background-estimation disc; not an area cutoff. |
| Fine sigma (px) | DoG | 1 | Small Gaussian scale. |

| | | | |
|---|---|---|---|
| Coarse sigma (px) | DoG | 8 | Large Gaussian scale (capped at 1/6 of the smaller image side; must exceed fine sigma). |
| Global floor multiplier | DoG | 1.3 | Object mean ≥ image mean + multiplier x SD. |
| Maximum eccentricity | DoG | 0.90 | Rejects elongated objects (0-1). |
| Local contrast z minimum | DoG | 1 | Minimum object contrast over local background; scaled by noise calibration when enabled. |
| Minimum shared pixels | Overlap | 3 | Pair qualification threshold. |
| Batch-noise scaling | DoG | On | Scales the z minimum by sample/reference noise. |

Defaults are starting points, not validated cutoffs for every microscope, objective or pixel size. Area limits and sigmas are in pixels and should be re-evaluated if the pixel size changes. Report all final values.

##### Validation rules enforced by the application

All parameters must be finite; minimum area ≥ 1 and maximum area ≥ minimum; thresholds non-negative; brush size positive; eccentricity between 0 and 1; coarse sigma greater than a positive fine sigma (DoG); minimum shared pixels a positive whole number.

#### 16. Reproducibility and reporting guidance

##### 16.1 What to record

- GGProfiler version, R version and package versions (EBImage, shiny, ggplot2, DT; the RDS snapshot stores sessionInfo()).
- Microscope, objective, pixel size, z-handling (single plane or projection method), and acquisition settings.
- Cropping criterion and the crop log.
- For each channel: segmentation method and all parameters; whether batch-noise scaling was used; calibration reference values and the images they were calculated from.
- Minimum shared pixels for overlap.
- Number of images processed, failed and zero-object, with reasons for any exclusion.
- Statistical test, correction, alpha and the unit of replication.

##### 16.2 Suggested Methods text (template)

Single-nucleus z-planes [or projections] were cropped in GGProfiler (v1.0) and paired by nucleus ID. P granules (PGL-1:mCardinal) were segmented by [top-hat / DoG] filtering and Mutator foci (MUT-16:GFP) by difference-of-Gaussians filtering (fine sigma [1] px, coarse sigma [8] px). Candidate objects were thresholded at the mean plus [multiplier] standard deviations of the positive filter response and retained if their area was [min]-[max] px, their mean intensity exceeded the image mean by [1.3] SD, their eccentricity was at most [0.90], and their local contrast z (object mean minus the median of a 4-px surrounding ring, divided by 1.4826 times the ring MAD) was at least [z_min], scaled by the ratio of the sample's median ring MAD to a reference value derived from [dataset]. For each nucleus, granule number, median size, median eccentricity, nearest-neighbor and radial spacing variability, dilation-based inter-channel apposition (7-px disc, chance-corrected) and size ratio were computed. All distances and areas are in pixels [convert with X um/px]. Images shown in figures were adjusted for brightness and contrast for display only; quantification used unadjusted images.

##### 16.3 Good practice

- Fix parameters on a tuning subset, then apply them unchanged to all conditions and cohorts.
- Acquire conditions that will be compared in the same session, with the same settings, and randomize or blind the order where possible.
- Include nuclei from the full age range during QC so that parameters do not favor one age.
- Analyze nuclei nested within animals for inference (Section 10).

#### 17. Limitations and interpretation cautions

- Spatial measurements are 2D and in pixels; they are not 3D distances.
- No nuclear segmentation: the "radial" measure uses the image center, so results depend on centered crops of similar size. Crop size and the position of the nucleus in the crop influence _radcv.
- _ring reports nearest-neighbor spacing variability, not ring structure.
- Peak-to-peak distances differ from edge-to-edge gaps.
- Proximity measures depend on the 7-px dilation and on a tissue domain that includes all configured channels; interpret them as relative, not absolute, associations.
- Local background rings are not corrected for neighboring objects, which can lower contrast in crowded nuclei.
- Noise scaling is a legacy heuristic whose appropriateness must be validated for each dataset.
- Median-based size and elongation summaries ignore within-nucleus heterogeneity; consult the object table when needed.
- Overlap of masks is not evidence of molecular interaction; check object counts alongside percentages.
- The Compare tab treats nuclei as independent (Section 10).
- Display contrast in overlays and figures does not reflect quantitative intensity.

#### 18. Troubleshooting

| Problem | Likely cause and remedy |
|---|---|
| "Enter every channel name" / "Channel names must also be unique." | Empty or duplicate names, or names that collapse after make.names (for example "A-1" and "A.1"). Rename. |
| "Remove all conditions before changing channel setup" | Setup was edited after adding conditions. Remove conditions, apply setup, add them again. |
| Files listed as excluded or invalid | Missing channel file, duplicate numeric ID, or no trailing number. Fix names and re-upload the full set under each marker. |
| "Calibrate the current dataset/settings first" | Noise scaling is on and calibration is absent or stale. Click Calibrate, or disable scaling (not recommended for comparative studies). |

| | |
|---|---|
| "Channel dimensions differ" | Channels of a nucleus have different pixel sizes. Re-crop all channels with the same rectangle. |
| "Unsupported image dimensions/ type" | Z-stack or unsupported file. Provide a single 2D plane or RGB image. |
| Many NA values | Too few objects (< 3 or < 4 for spacing measures) or one channel empty. Check segmentation in QC. |
| Comparison returns errors | Fewer than three finite observations per condition, or constant data. |
| Results disappeared | Settings, uploads or setup changed after extraction; rerun extraction. |
| Folder saving of crops fails | Application is hosted on a server or the folder is not writable. Use the ZIP download. |
| Font warning in PNG export | Helvetica is not available to the device. Use PDF export or install a suitable font. |
